# Flow similarity model predicts the allometry and allometric covariation of petiole dimensions

**DOI:** 10.1101/2023.01.05.522893

**Authors:** Charles A. Price

**Affiliations:** National Institute for Mathematical and Biological Synthesis, University of Tennessee, Knoxville, TN, 37996-3140, USA; School of Biological Sciences, University of Western Australia, Perth, Western Australia 6009, Australia

## Abstract

Allometric relationships for plants, plant organs and plant parts, have long generated interest among biologists. Several prominent theoretical models based on biomechanical and/or hydraulic arguments, have been introduced with mixed support. Here I test a more recent offering, flow similarity, which is based on the conservation of volumetric flow rate and velocity. Using dimensional data for 935 petioles from 43 angiosperm species, I show that both the intraspecific and interspecific petiole allometries are more closely aligned with the predictions of the flow similarity model than that of elastic or geometric similarity. Further, allometric covariation among empirical scaling exponents falls along predicted functions with clustering around the flow similarity predictions. This work adds to the body of literature highlighting the importance of hydraulics in understanding the physiological basis of plant allometries, identifies previously unknown central tendencies in petiole allometry, and helps to delineate the scope within which the flow similarity model may be applicable.

## INTRODUCTION

The study of morphology has a venerable history in biology, with attempts to model biological form going as far back as Leonardo DaVinci (Richter, 1970) and Galileo Galilei (Galilei, 1638) and championed early in the last century by luminaries such as D’Arcy Thompson and Julian Huxley (Thompson, 1917; Huxley, 1932). This area has seen great interest of late with several prominent models having been published that invoke physical processes to predict the scaling of biological form (West et al., 1999; Banavar et al., 2002; Savage et al., 2010; Banavar et al., 2014).

For plants, modelling attempts typically invoke hydraulic processes (Shinozaki et al., 1964; Niklas and Spatz, 2004), biomechanical processes (McMahon and Kronauer, 1976), or combinations of the two (West et al., 1999; Savage et al., 2010). A common feature of many of these approaches is that they predict branches of differing size will exhibit self-similarity in their dimensions due to underlying biomechanical or hydraulic constraints. For example, the elastic similarity model (McMahon, 1973; McMahon and Kronauer, 1976) predicts the maximum height to which an idealized column can extend before it buckles due to self-loading, which depends on the column radius and material properties density and elastic modulus. The model also predicts that the ratio of a branch’s length to its deflection due to self-loading, will be constant across branches of different size. A common null model in biology is that of geometric similarity which predicts that biological objects will have allometric exponents similar to those expected for self-similar geometric shapes such as spheres (Galilei, 1638; Niklas, 1994). A recent effort that has garnered considerable attention is West, Brown and Enquist’s fractal similarity model, which assumes elastic similarity, but goes on to make a suite of additional predictions including the scaling of leaf number and plant mass (West et al., 1997, 1999; Enquist and Niklas, 2001; Enquist and Niklas, 2002). This body of work has catalysed great interest in the origin of allometric scaling relationships, but many have questioned the models scope and mechanistic underpinnings for plants (Dodds et al., 2001; Niklas, 2004; Muller-Landau et al., 2006; Petit and Anfodillo, 2009; Price et al., 2010; Price et al., 2012).

Recently, Price et al.(2022), introduced a “flow similarity” model which similarly predicts scaling relationships between different dimensions (length (*l*), diameter (*d*), surface area (*SA*), and volume (*V*)) of branching networks within and across plants. Price et al.(2022) showed that several parts of plant branching networks including juvenile tree stems, interspecific terminal branches, and interspecific petioles have allometric relationships that are generally consistent with the predictions of the flow similarity model (detailed derivations and explanation can be found in Price et al., 2022). The model is based on the Hagen-Poiseuille equation for laminar flow in a cylindrical tube and relies on two principal model assumptions/constraints, velocity preservation and the conservation of volumetric flow rate (hence “flow similarity”) which together predict a constant pressure drop (note that any two of these constraints predicts the third). Two additional assumptions, a constant bulk tissue density and an isometric relationship between internal (xylem) and external branch numbers are invoked (exceptions to these assumptions are considered in the manuscript) to yield a series of predictions including *l* ∝ *r* ^2^ which leads to *SA* ∝ *V* ^3/4^ scaling, and the four additional allometric relationships described in Table 1. Price et al. (2022) showed that interspecific allometric relationships for leaf petioles were better fit by the flow similarity model than elastic similarity, but did not consider intraspecific petiole allometries, or allometric covariation among intraspecific exponents as is done here.

**Table 1.**
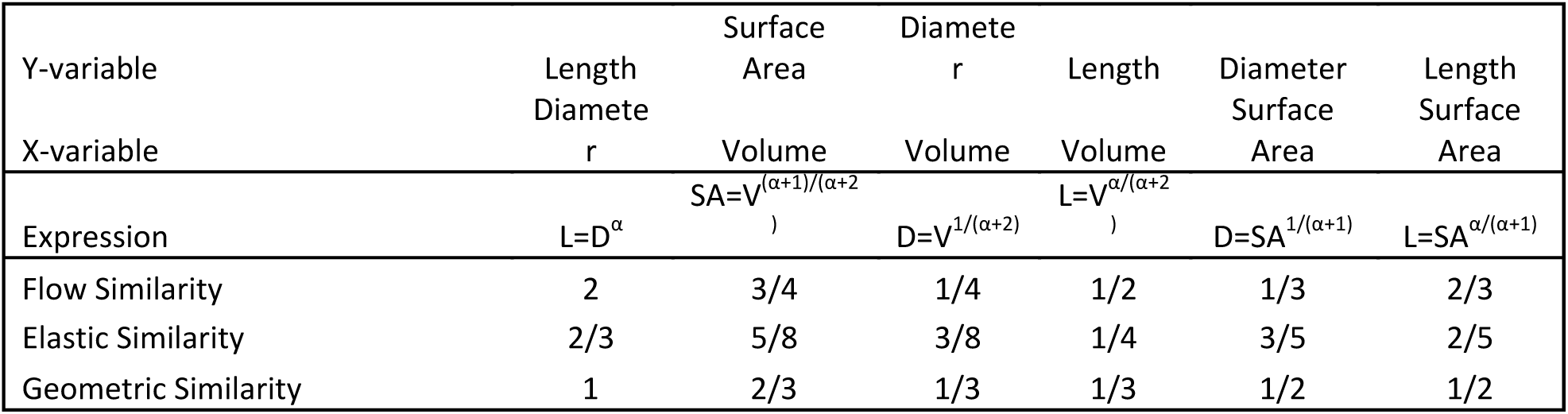
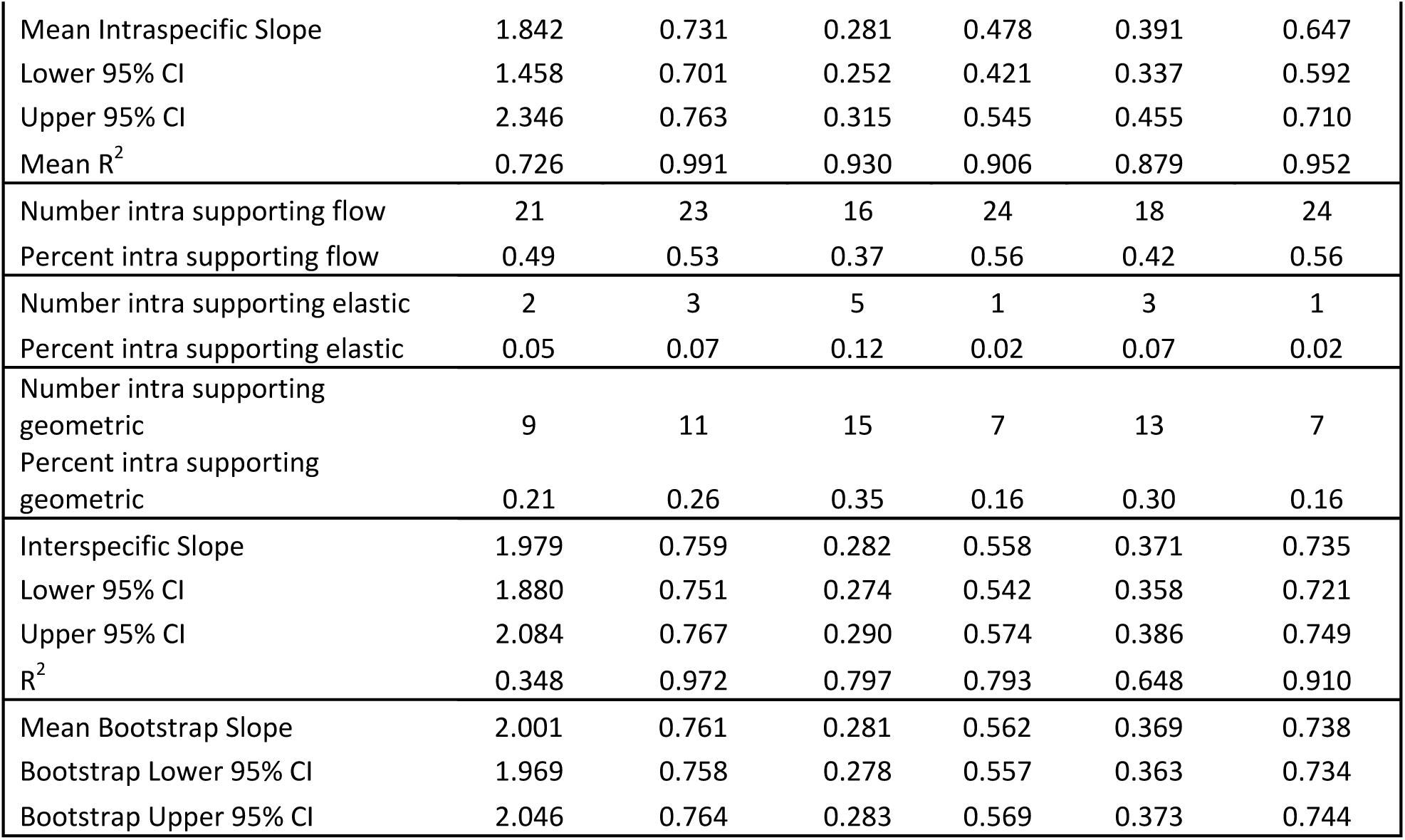
Dimensional variables, model predictions, and empirical results. The first two rows represent the dimensional variables in question, and row three the expression from the model for the relationship between the two. Rows 4-6 are the predictions from the three models evaluated (flow similarity when α=2). Rows 7-10 represent intraspecific results, and rows 11-16 the number and percent of the intraspecific results supporting each model. Rows 17-20 are the interspecific slope results, with 21-23 the bootstrap results for same.

Leaves and supporting petioles typically develop from primordia below the shoot apical meristem and undergo numerous rounds of cell division and enlargement before reaching full expansion. Growing petioles serve two primary roles, supporting the mass of leaf tissue and delivering resources to and from leaves. Most studies of petiole biomechanics to date have been either ecological, i.e., how resource or stress gradients influence petiole form and function (Niklas, 1996; Anten et al., 2010), or have compared the mechanical behaviour of basic petiole types or shapes (Etnier and Villani, 2007). For example Niklas showed that the leaves of simple and palmately compound species behave effectively as non-tapered, cantilevered columns and behave differently than pinnately compound leaves (Niklas, 1991). Vogel examined the resistance to twisting and bending in the leaves of several species and showed that non-circular petioles tend to be more flexible in twisting than circular ones (Vogel, 1992).

Despite interest in this area, many basic questions remain regarding the geometry and design of leaf petioles. Do the petioles from different species have similar geometries? For example, if we examine the allometry of leaf petioles over the course of leaf expansion within and across species, do different species have similar scaling exponents or is there wide variation in the allometry of petiole morphology? Looking across species, do we see systematic covariation of these intraspecific scaling exponents? Further, are the central tendencies for collections of empirical intraspecific or interspecific allometric exponents consistent with the expectations for any of the aforementioned scaling models?

Here I address these and other questions by examining intraspecific allometric relationships among the linear dimensions of petioles of 43 temperate angiosperm species. Using the method of multiple working hypotheses (Chamberlain, 1897; Platt, 1964), I show that the flow similarity model shows greater agreement with the allometric data at both inter and intraspecific levels than either elastic or geometric similarity. I also show that covariation among allometric exponents is in strong agreement with model predictions (Price and Weitz, 2011). The implications of this work for scaling in plants, and the potential broader ecological and evolutionary contexts are discussed.

## MATERIALS AND METHODS

935 leaves from 43 species for a mean of ∼22 individuals per species, were collected during the summer of 2007 within the greater Atlanta region (Lat/Long 33 75 – 84 38). Data were collected initially as part of a study on the allometry of leaf surface area (Price et al., 2009), but only petiole diameters were used; petiole length, surface area and volume were not reported nor interpreted. Species were chosen based on local availability. Initially we had hoped to analyse differences between simple and compound leaves but following data collection there were not enough compound leaved species for a meaningful statistical comparison, thus we lump both types together here. The petioles of all species were approximately cylindrical in nature.

For each fresh leaf, petiole length, including the rachis in six compound leaved species, was measured with a ruler, and petiole diameter (just beyond any existing pulvinus) was measured twice at the base with digital callipers. For each species, as large a range of leaf and petiole sizes as could be found was collected, with representative sampling across the size range. Each petiole was assumed to have a cylindrical shape, and surface area and volume were estimated from the length and diameter measures using standard geometric formulas.

Bivariate relationship among the four dimensional variables (*l, d, SA, V*) were estimated using standardized major axis regression (SMA). Regression exponents were estimated using the software package SMATR (Falster et al., 2003). SMA regression is typically used in allometric studies where both variables have associated measurement error and one wants to test if a slope equals a certain value, as is the case here (Warton et al., 2006). All data were log transformed before regression fitting to meet the regression assumption of homogeneity of variance. Regression functions were fit to individual species data (intraspecific) and to all data together (interspecific). As there are multiple comparisons for each intraspecific model test, I utilize a Holm-Bonferroni correction for family wise error rate (Holm, 1979; Groppe, 2022) and report the corrected number of significance tests for each of the three models evaluated.

For interspecific regressions, species were represented by different numbers of individuals ranging from 18 to 40 leaves per species. To ensure that differences in sample size didn’t influence interspecific slope estimates, I employed a bootstrapping approach. 18 individuals were drawn at random from each species, and a SMA regression line was fit to each interspecific relationship. This procedure was repeated 10,000 times to yield a distribution of slope estimates from which I could calculate a mean interspecific slope and 95% confidence intervals.

Among the four dimensional variables measured or estimated here (*n*), length, diameter, surface area and volume, per the binomial coefficient (*n*!/((*n*–*k*)! *k*!) there are six possible pairwise (*k*) combinations. Similarly, there are 15 pairwise relationships among the six allometric exponents. Previous work (Price et al., 2022) has shown that if one represents the length vs diameter allometric relationship as *L* ∝ *D*^*α*^, where α represents the scaling exponent, one can predict covariation functions as a function α, among all of the aforementioned 15 pairwise allometric exponent relationships.

I evaluated these predicted covariation functions by plotting the observed allometric exponents and predicted functions together. The fit of the predicted function to data was assessed by the coefficient of determination (*R*^2^) using the standard formula *R*^2^=1-(*SS*_*Res*_/*SS*_*Tot*_). SS_Res_ refers to the residual sum of squares and is given by *SS*_*Res*_ = ∑(*y*_*i*_ − *f*_*i*_)^2^, where *y*_*i*_ corresponds to each observed value and *f*_*i*_ corresponds to the predicted value from the covariation function. SSTot refers to the sum of squares total and is given by 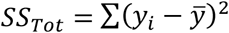, where 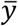 refers to the mean observed y value.

## RESULTS

The six intraspecific allometric relationships across the four dimensional variables typically had high coefficients of determination (*R*^2^) (Fig. 1, Table 1). The mean *R*^2^ over all 258 bivariate relationships examined was 0.9 (Table 1). The weakest relationship was length vs diameter with a mean intraspecific *R*^2^ of 0.73, and the strongest relationship was surface area vs volume with a mean *R*^2^ of 0.99 (which isn’t surprising as both surface area and volume are estimated from length and diameter).

**Figure 1.**
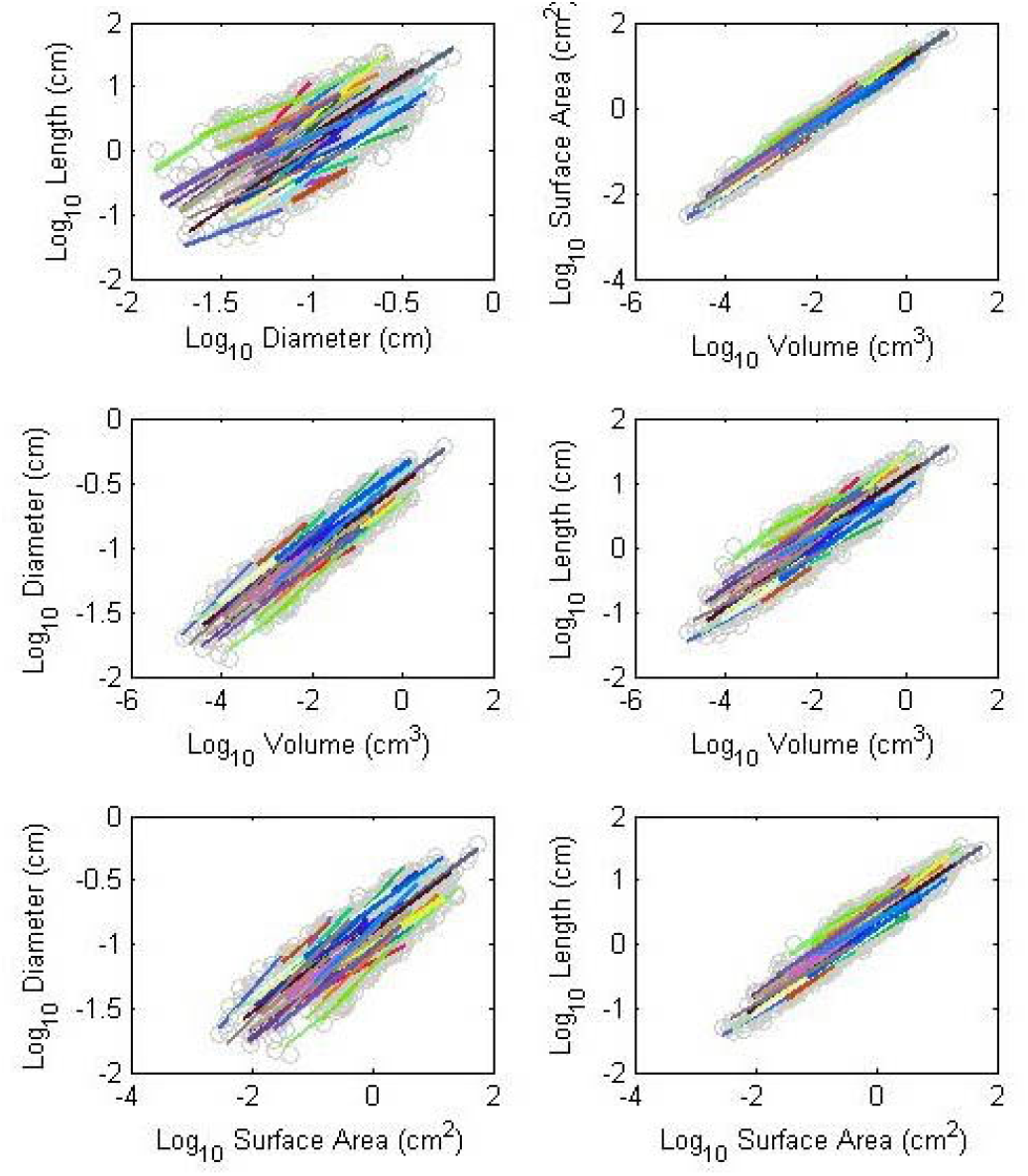
Bivariate relationships between the four, dimensional variables considered, petiole length, basal diameter, surface area and volume. Colored lines represent SMA fits to individual species data and are included to help visualize the ordinal spread (Panel A), or lack thereof (Panel B) in each plot. Note that despite the fact the for a given petiole diameter, there exists close to two orders of magnitude of variability in petiole length, yet all species collapse around what appears to be a single surface area to volume function.

Out of 258 pairwise relationships, 185 (∼72%) had slope estimates who’s 95% confidence intervals included the prediction from the flow similarity model (Table 1) following the Holm-Bonferroni correction. By comparison, 127 out of 258 (∼49%) and 35 out of 258 (∼14%) relationships had confidence intervals that included the geometric similarity and elastic similarity model predictions, respectively (Table 1).

Interspecific allometric relationships had similar slopes to the mean intraspecific values, with higher variance as would be expected (Table 1). Bootstrap slope estimates were very close to those obtained without bootstrapping, thus differences in sample size had little influence on interspecific slope estimates here.

The distribution of slopes in Panels A-F (Fig. 2) are all modal and while no normality tests were performed (such tests are notoriously sensitive to single outliers and small sample sizes), all distributions look fairly normal. Species with higher correlation coefficients (*R*^2^), tended to cluster more closely to model predictions (Fig. 3).

**Figure 2.**
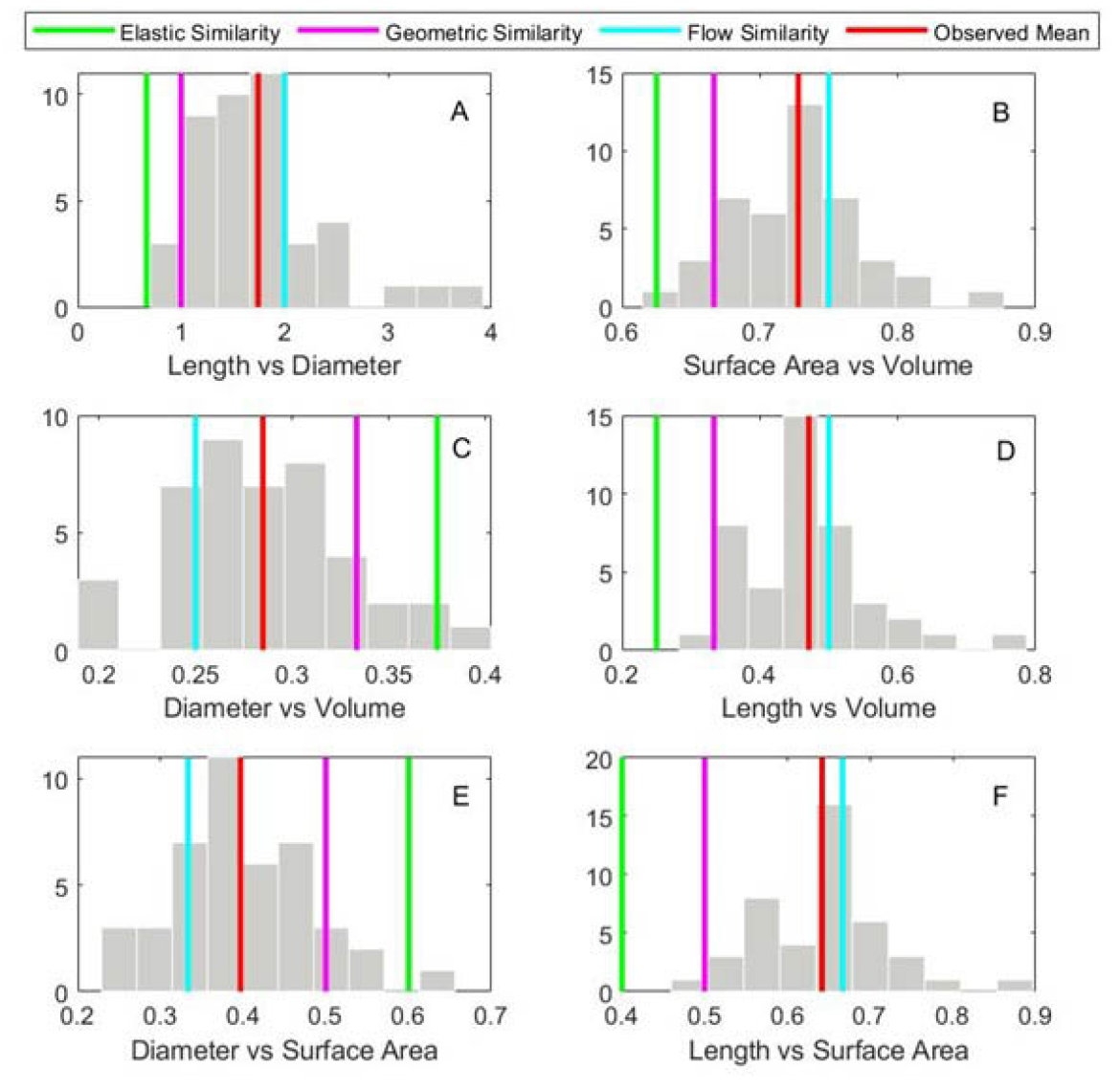
Frequency distributions for the slopes of the six intraspecific allometric relationships. All distributions are approximately model and fairly symmetric. Note the mean value for each distribution (red line) is closest to the flow similarity model expectation in all cases.

**Figure 3.**
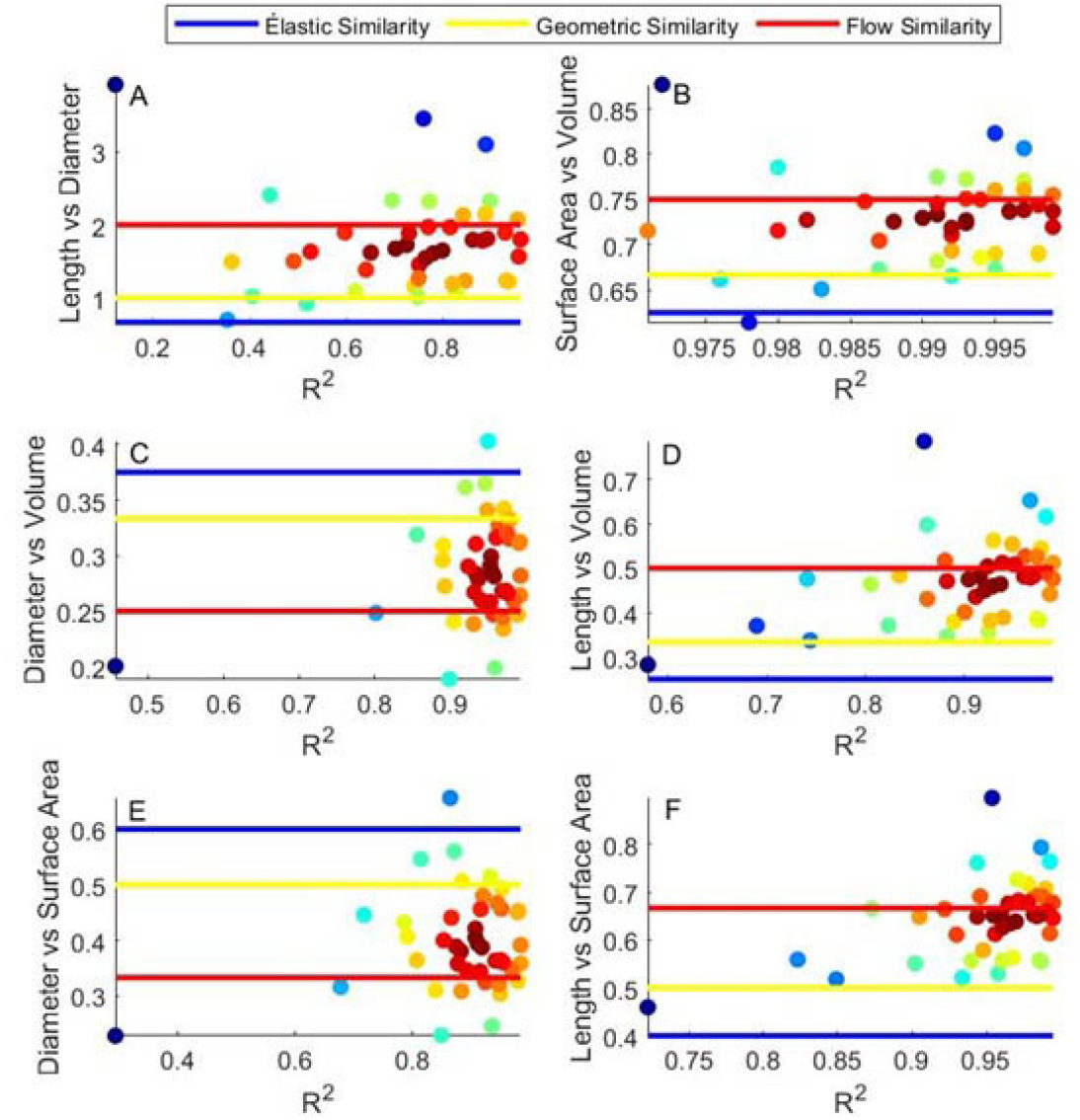
Bivariate plots for the six allometric relationships considered. Each observed allometric exponent is plotted as a function of its corresponding *R*^2^ value for each relationship. Point colors correspond to the mean distance to all other points and thus indicate neighbourhood density. Note that as the *R*^2^ values increase, the points tend to converge at or near the expectation for the flow similarity model although for Panels C and E there is also a modest clustering near the expectation for the geometric similarity model as well.

The agreement between predicted covariation functions and observed allometric exponents is quite good. As is seen in Fig. 4, the predicted function has the same curvature as the data in all cases. The amount of variation in the data explained by the predicted function (*R*^2^) is generally high, with a minimum value of 0.49 (Fig. 4, Panel K), and a maximum value of 0.99 (Fig. 4, Panels I and N). The mean *R*^2^ value across all 15 functions is 0.85.

**Figure 4.**
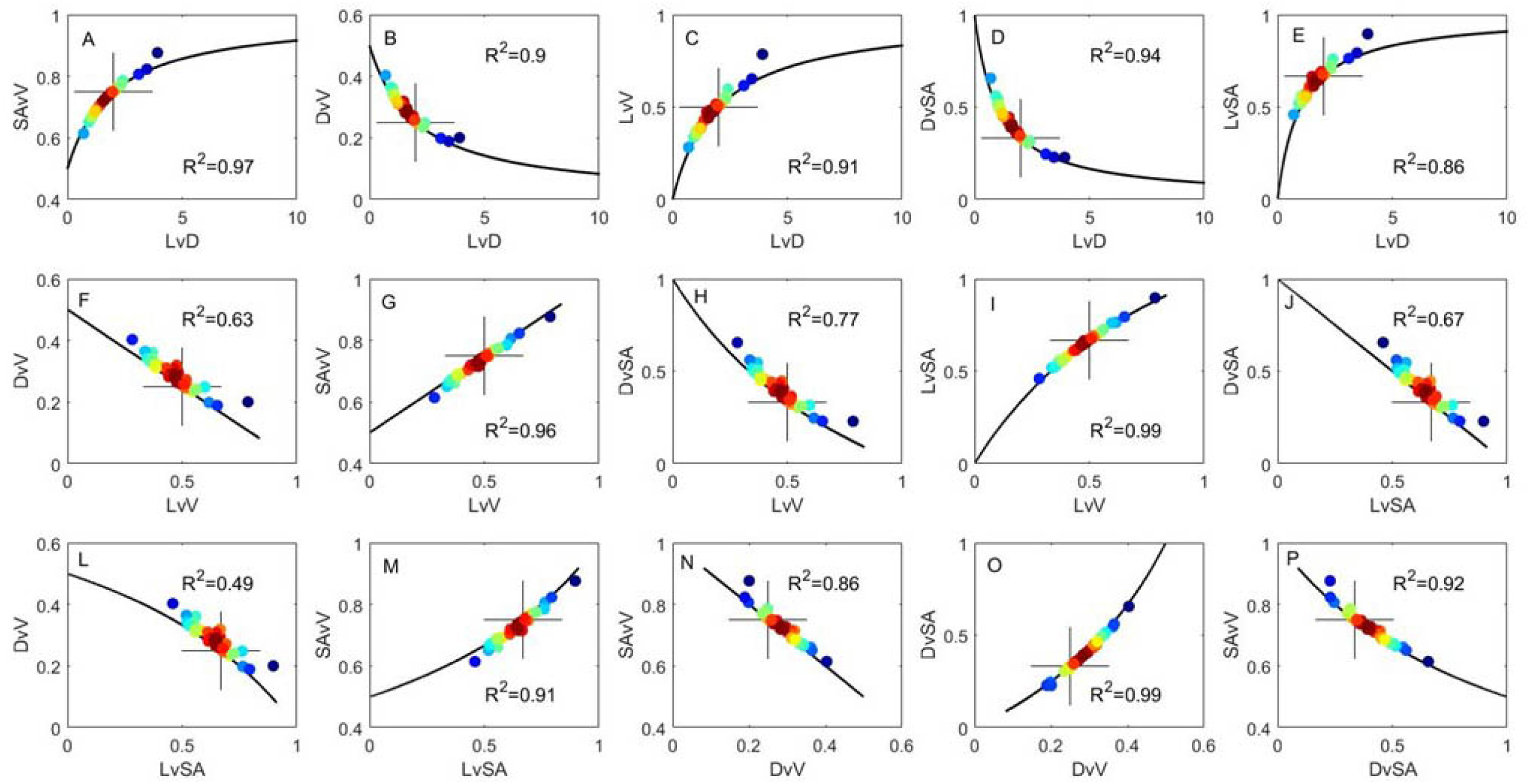
Predicted covariation functions for the 15 possible pairwise relationships among allometric exponents with the prediction from the flow similarity model (when α=2, Table 1) represented by a black cross. The blue line in each panel represents the predicted covariation function (Table S1), the points represent observed combinations of exponents for the 43 species. Point colors correspond to the mean distance to all other points and thus indicate neighbourhood density. *R*^2^ values for each covariation relationship are shown in each panel. Note the fairly strong agreement between model predictions and observed data.

## DISCUSSION

Allometric relationships among petiole dimensions are generally quite strong. As seen in Fig. 1 and Table 1, the slopes are all fairly similar to one another and typically have high *R*^2^ values. While Panels A and C-F in Fig. 1 all have around one to two orders of magnitude variability in the Y-variable for any given X-variable, for the relationship between surface area and volume (Panel B), all data collapse substantially. One might expect the relationship between surface area and volume to be constrained as both are estimated as a function of length and diameter. However, there is no a priori theoretical reason based on geometry alone why slopes would converge around a single value (save geometric similarity which assumes a self-similar shape). The smallest surface area one could have for a given volume is of course a sphere, but petioles are obviously better approximated by cylinders. While biologically unrealistic, one could have an extremely thin cylinder with minimal volume and very large surface area, thus from a strictly theoretical standpoint, there is a lower (2/3), but no upper bound for the slope values in Fig. 1B.

Generally, there is very good agreement between the mean intraspecific slope and the predicted value from the flow similarity model. In contrast, there is less agreement with the elastic or geometric similarity models, with geometric similarity outperforming elastic similarity (Table 1 and Fig. 2). Moreover, as the *R*^2^ values for each relationship increase, they appear to converge on values close to those predicted by flow similarity (Fig. 3). This aspect of allometric data is often ignored, and suggests that some of the slopes that are different than expected may be due to poor correlations, and perhaps worth re-examining in those species.

As seen in Fig. 4 there is strong agreement between predicted covariation functions and observed data. This is perhaps not surprising as the predicted functions each form a continuum that represents the expectation for all possible series of self-similar cylinders, specifically all possible values of α (Table S1). The model-data agreement tells us not only that many petioles may be well approximated by self-similar cylinders, but also helps to delineate what parts of the theoretical morphospace are occupied, and which are not. For example, no length vs diameter exponents exceeded a value of 4. While larger data collections might better delineate the range of this and the other exponents, it is clear that we are unlikely to find values on the order of 10 or 100. By evaluating mechanistic models such as those discussed here (flow, elastic, or geometric similarity) we can begin to question what constrains petioles to be within this range of the morphospace and not another.

Another benefit to delineating the theoretical covariation space, is that one can begin to ask questions with a more ecological or evolutionary focus. For example, if one were to examine the petiole or branch allometry of a group of species from a single genus or family, one could begin to examine how shifts along these interrelated covariation functions corresponded to adaptation within the group. As another example, one might envision a study looking at the mean exponent values for groups of species found in different environments to see if there are systematic shifts along these functions associated with variability along resource or stress gradients.

The poorer performance of the geometric similarity model simply indicates that strictly proportional relationships between the petiole dimensions are not maintained as petioles get larger. The poor performance of the elastic similarity model suggests that its underlying mechanisms may not have a strong influence on petiole form, or at least that other constraints or selective pressures play a stronger role in shaping petiole allometry. It is not currently known whether biomechanical or hydraulic drivers have a stronger influence on petiole form in this context. The flow similarity approach is based primarily on hydraulics and suggests that biomechanics become more important as plant get large and the influence of self-loading increases (Price et al., 2022). This study helps to delineate the scope of the model as the data do seem to support its applicability for petioles, the allometry of which may be more influenced by hydraulics.

The flow similarity model was developed for bifurcating networks, which can include petioles. However as discussed in Price et al. (2022), selection may also be acting on hydraulic traits that reflect the species level hydraulic constraints.. Specifically, it has been shown that branching that follows *l* ∝ *r* ^2^ scaling preserves sapwood specific conductivity (*K*_*S*_), defined as 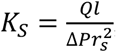, where *r*_*s*_ is the sapwood radius. Conservation of sapwood specific conductivity has been shown in terminal branches of trees (McDowell et al., 2002; Sellin et al., 2008), but to my knowledge has not been measured intraspecifically in petioles.

The relatively good agreement between the flow similarity predictions and data provides tentative support for the underlying modelling approach. Combined with the results in Price et al. (2022) for terminal stems and saplings, the work presented here helps to delineate the scope of the flow similarity model. However, as recently discussed in the context of testing metabolic scaling theory (Price et al., 2012), subsequent tests of the simplifying assumptions and modelling constraints are required to determine what mechanisms ultimately drive branch and petiole form. With respect to the assumption of a constant bulk tissue density, it seems reasonable to assume that within a species, petiole tissue density would not vary much, but empirical support for this would be useful. Further, studies examining petiole specific conductivity, in those species in which is its empirically tractable, would help to determine the extent to which this is a species-specific trait. Additional tests examining flow velocities and pressure changes within and across petioles of differing sizes might help delineate if the underlying mechanism is consistent with those the model invokes.

As discussed in Price et al. (2022), the simplifying assumptions of the flow similarity model are most closely met when the dimensions of the plant parts in question (stems, petioles, veins) are governed more by hydraulics than biomechanical demands as might be observed in small plants or the distal portions of large plants. As plants increase in size, the demands of self-loading will likely drive scaling relationships towards elastic similarity and scaling relationship for large size ranges and data collections may become non-linear with slope estimates falling between flow similarity and elastic similarity predictions (see discussion in Price et al. 2022).

## CONCLUSIONS

Scaling hypotheses based on physical first principles have a long history in biology (Niklas, 1994). The flow similarity model adds to this body of work by considering the allometric relationships that might emerge under relatively simple, and arguably parsimonious model constraints, the conservation of volume flow rate and velocity. The data presented herein demonstrate that petiole allometries are more consistent with the flow similarity model than other prominent scaling models. However, the model is not intended to be universal, and violations of its simplifying assumptions are expected, particularly as trees or leaves become large. Thus, subsequent tests that help to delineate to scope within which the model might help us understand the scaling of plant form, are of value in determining it usefulness.

## ACKNOWLEDGEMENTS

CAP wishes to acknowledge the support of an Australian Research Council (ARC) Discovery Early Career Researcher Award (DECRA - DE120101562) and a fellowship from the National Institute for Mathematical and Biological Synthesis (NIMBioS, USA).

## AUTHOR CONTRIBUTIONS

CAP designed the study, collected the data, performed the analysis, and wrote the manuscript.

## DATA AVAILABILITY

The raw data for the species names and petiole dimensions are available on Zenodo.

